# A fast-acting inhibitor of blood-stage *P. falciparum* with mechanism distinct from artemisinin and chloroquine

**DOI:** 10.1101/2024.08.12.607553

**Authors:** Stephanie Kabeche, Thomas Meister, Ellen Yeh

## Abstract

Artemisinins are first-line treatment for malaria, prized for their extremely fast reduction of parasite load in patients. New fast-acting antimalarial compounds are urgently needed to counter artemisinin resistance, but the fast parasite reduction observed with artemisinins is rare among antimalarial compounds. Here we show that MMV1580853 has a very fast *in vitro* killing rate, comparable to that of dihydroartemisinin. Near-complete parasite growth inhibition was observed within 1 hour of treatment with MMV1580853 and dihydroartemisinin, while chloroquine, another fast-acting antimalarial, showed partial growth inhibition after 1h. MMV1580853 was reported to inhibit prenyltransferases, but its fast killing rate is inconsistent with this mechanism-of-action and we were unable to validate any of 3 annotated *P. falciparum* prenyltransferases as MMV1580853 targets. MMV1580853 also did not phenocopy the inhibition phenotype of either chloroquine or dihydroartemisinin. These results indicate that MMV1580853 has a distinct mechanism-of-action leading to a very fast killing rate. MMV1580853 compound development and investigation of its mechanism-of-action will be critical avenues in the search for drugs matching the remarkable clinical efficacy of artemisinin.

---

The development and spread of artemisinin resistance in *Plasmodium falciparum* threatens malaria eradication efforts. Artemisinin and its derivatives (ARTs) are the fast-acting compounds in frontline combination treatments for acute malaria. There are currently no alternatives to ARTs available. Thus there is an urgent need to develop new antimalarials with new modes-of-action that are not cross-resistant, yet preserve the fast action of ARTs [1].

We previously tested 400 compounds from the open-access Pandemic Response Box released by Medicines for Malaria Venture for growth inhibition of *P. falciparum* blood cultures [2,3]. One compound, MMV1580853, showed potent inhibition of blood-stage *Plasmodium falciparum* W2 growth with EC_50_= 13.0 nM (12.5-13.4 nM) (Fig S1). To determine if MMV1580853 is fast-acting, we performed the *in vitro* parasite viability fast assay which differentiates the parasite killing kinetics of antimalarial compounds compared to control drugs as slow (atovaquone), medium (pyrimethamine), or fast (chloroquine) [4]. MMV1580853 caused >99% parasite growth inhibition within the first 24h of treatment, similar to chloroquine, categorizing it as fast-acting (Fig 1A). More sensitive assays are needed to differentiate between fast-acting compounds identified in this assay. To further characterize the fast action of MMV1580853, we tested 1h drug pulses in ring-stage parasites, an assay designed to mimic clinical drug exposure in patients and that distinguishes fast-acting antimalarials chloroquine and artemisinin [5]. Surprisingly, MMV1580853 demonstrated near-complete parasite growth inhibition after just 1h of treatment, comparable to treatment for the full parasite replication cycle (Fig 1C). Compared to known fast-acting antimalarials, only dihydroartemisinin (DHA) showed similarly fast action within 1h, whereas chloroquine showed partial growth inhibition in 1h. As *in vitro* killing rates of antimalarial compounds correlate with *in vivo* parasite clearance [4], these results indicate that MMV1580853 derivatives may have clinical utility as substitutes for current fast-acting antimalarials.

**Figure 1.**
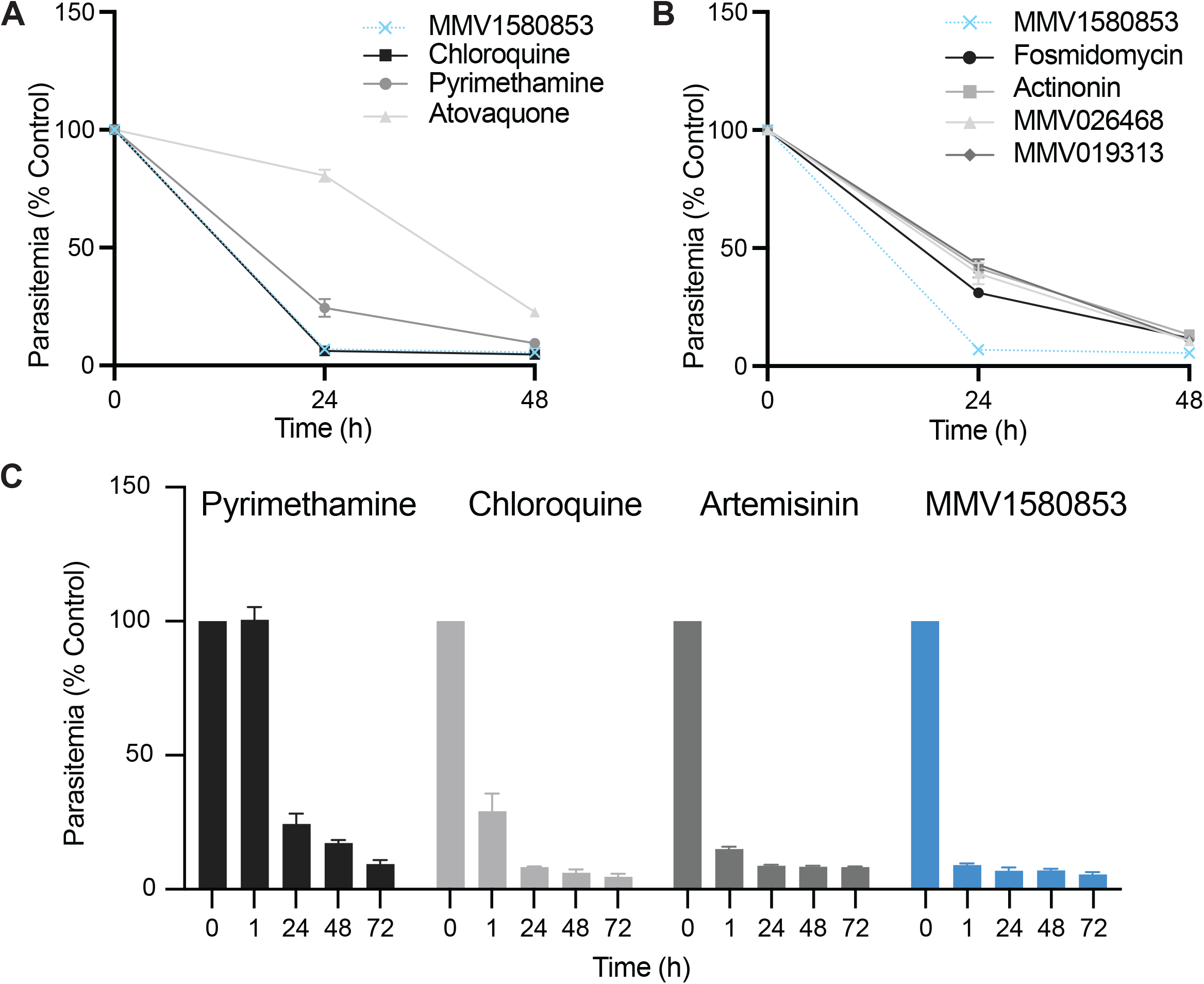
MMV1580853 is a fast-acting antimalarial inhibitor. A-B. Parasite viability assay to distinguish slow, moderate, and fast acting antimalarials. MMV1580853 treatment is compared to control compounds (A) and known isoprenoid and apicoplast inhibitors (B). C. Ring-stage viability after 1h drug pulse compared to full growth inhibition at 24-72h time points. Results are the mean and 95% CI of three independent experiments after subtraction of background fluorescence of uninfected RBCs. Growth is normalized to untreated control.

MMV1580853 has been reported to inhibit the *in vitro* activity of human farnesyl pyrophosphate synthase (FPPS; IC_50_= 1.8μM) [6] and bacterial undecaprenyl diphosphate synthase (UPPS; IC_50_= 0.1μM) [7], both prenyltransferases in isoprenoid pathways. In *Plasmodium*, isoprenoid precursor biosynthesis occurs in the plastid organelle, the apicoplast, while later isoprenoid product biosynthesis by prenyltransferases occurs in the cytoplasm. *In vitro* killing rates are also indicative of the mechanism-of-action of antimalarials: compounds with shared mechanisms-of-action have similar killing rate profile and fast-acting compounds typically target functions in ring-stage parasites [8]. We measured the killing rates of known inhibitors of *Plasmodium* isoprenoid precursor biosynthesis (fosmidomycin) [9], prenyltransferase (MMV019313) [10], and apicoplast biosynthesis (actinonin) [11]. The selected isoprenoid and apicoplast inhibitors all showed a medium killing rate, consistent with the peak expression of isoprenoid and apicoplast genes at >30h post-infection in the 48h parasite life cycle (Fig 1B) [12].

Though the fast parasite killing kinetics of MMV019313 is inconsistent with a mechanism-of-action in isoprenoid pathways, we specifically tested whether MMV1580853 targets the 3 prenyltransferases annotated in the *P. falciparum* genome: farnesyl/geranylgeranyl diphosphate synthase (F/GGPPS; PF3D7_1128400), octaprenyl pyrophosphate synthase (OPPS; PF3D7_0202700), and cis-prenyltransferase (CPT; PF3D7_0826400). Conditional knockdown (KD) strains were generated by addition of a *C*-terminal 3xhemagglutinin (HA) tag and a 3’ UTR RNA aptamer sequence that binds a TetR-DOZI repressor to the endogenous gene locus (Fig S1) [13]. At ≥500 nM anhydrotetracycline (aTC), the aptamer is unbound and the targeted transcript is maximally translated. Absent aTC, repressor binding inhibits translation resulting in growth defects within the first replication cycle (Fig 2A). Partial translation repression at 5 nM aTC delays growth defects (Fig 2B) but sensitizes parasites to inhibition of the prenyltransferase during the first replication cycle compared to the unrepressed condition [11]. As a positive control, we compared the dose-response of *PfFPPS*/GGPPS KD strains at 5 and 500 nM aTC against its known inhibitor, MMV019313 [10,14], during the first replication cycle. We confirmed the leftward shift of the dose-response at 5 nM aTC compared to 500 nM aTC, indicating sensitization to the inhibitor targeting *Pf*FPPS/GGPPS (Fig 2C). In contrast, the dose-response to MMV158083 was unchanged at 5 and 500 nM aTC in *Pf*FPPS/GGPPS, *Pf*OPPS, and *Pf*CPT KD strains, indicating that these prenyltransferases are unlikely to be the targets of MMV1580853 (Fig 2D-F). Taken together, our results do not support a mechanism-of-action targeting isoprenoid pathways for MMV1580853.

**Figure 2.**
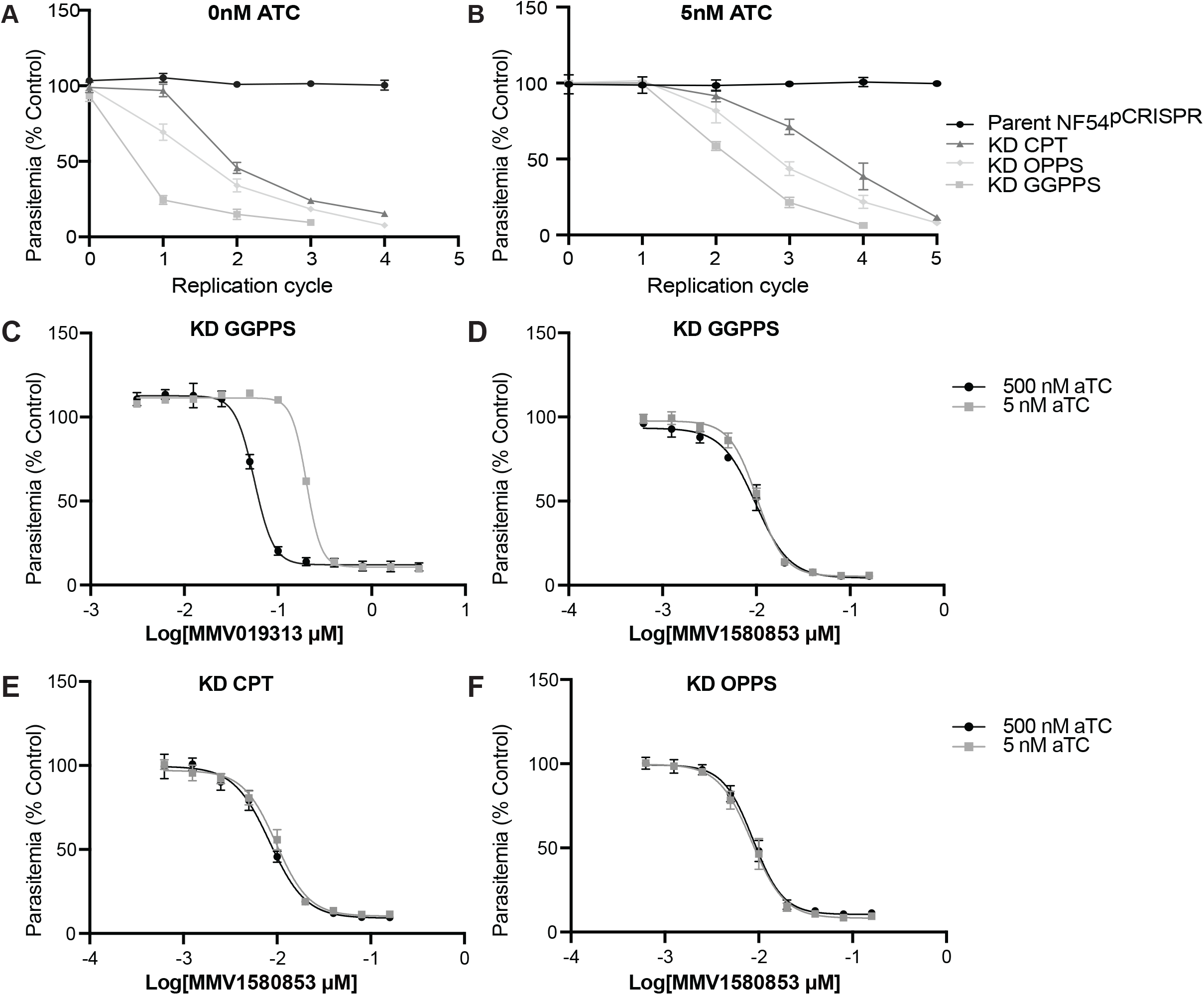
MMV1580853 does not inhibit known *P*.*falciparum* prenyltransferases. A-B. Time course of growth inhibition of *P. falciparum* KD strains at 0 nM (A) or 5 nM (B) aTC. C-F. Dose-dependent growth inhibition of *P. falciparum* in indicated KD strains at 5nM and 500nM aTC against MMV019313 (C) and MMV158083 (D-F). Results are the mean and 95% CI of 3 biological replicates. Growth is normalized to untreated control.

Since MMV1580853’s most distinctive phenotype was its fast action, we tested whether it shares a mechanism-of-action with clinically-used fast-acting antimalarials, chloroquine and artemisinin. Chloroquine blocks hemoglobin degradation in the parasite food vacuole causing distinctive morphologic changes and accumulation of hemoglobin [15]. Parasites were treated with either chloroquine, pyrimethamine, or MMV1580853, and their morphology were evaluated by Giemsa stain over the 48h replication cycle. As reported previously, chloroquine-treated parasites showed characteristic food vacuole swelling and pigment clumping (Fig 3A). In contrast, MMV1580853 showed stalled development but with no visible swelling or pigment clumping. Similarly, while hemoglobin accumulated under chloroquine treatment, this was not observed under MMV1580853 treatment (Fig 3B) [16,17].

**Figure 3.**
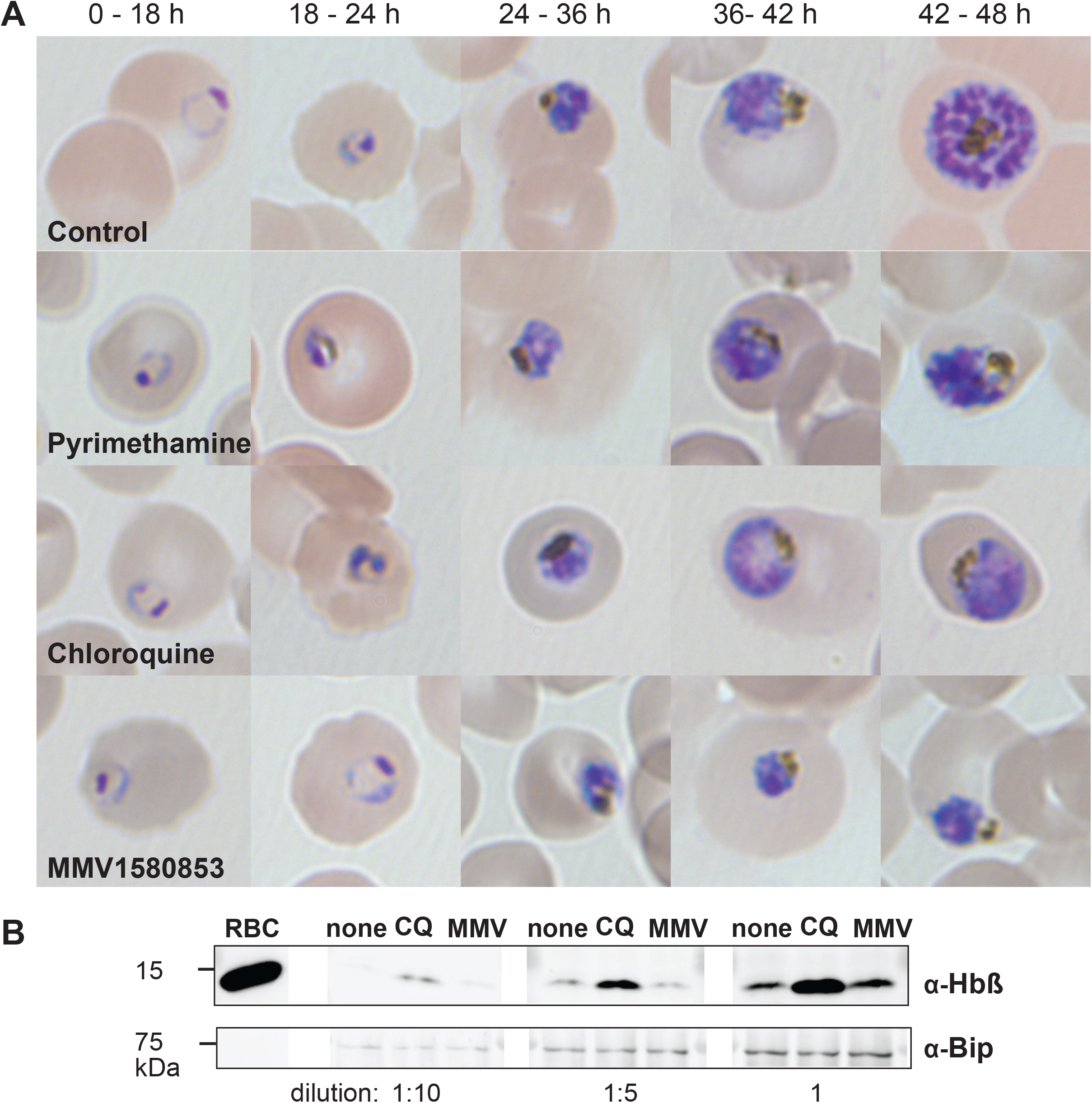
MMV1580853 treatment does not phenocopy chloroquine inhibition. A. Time course of food vacuole morphology defects in inhibitor-treated *P. falciparum* parasites evaluated by light microscopy B. Accumulation of undegraded hemoglobin in inhibitor-treated *P. falciparum* parasites by Western blot. Early trophozoite parasites (3D7) were grown for 10 h in the presence or absence of indicated drugs at concentrations 3x EC_50_. Protein was loaded at 1/10 (left set), 1/5 (middle set), and 1x (right set) dilutions. Abbreviations as follows: RBC, red blood cell; none, no drug; CQ, chloroquine; MMV, MMV1580853. Bip serves as a loading control.

Distinctive effects of artemisinin treatment have also been documented, including proteosome disruption resulting in accumulation of polyubiquitinated proteins and ER stress resulting in increased elF2α phosphorylation [18–20]. Consistent with these reports, we observed accumulation of ubiquinated proteins and increased elF2α phosphorylation with DHA treatment (Fig 4A-B). However, neither phenotypic effects were observed upon MMV1580853 treatment. Moreover, artemisinin-resistant strains, Dd2 Kelch13 C580Y and R539T, remained susceptible to MMV158053 with EC_50_= 0.0107 nM (0.0098-0.0117 nM) and 0.0114 nM (0.0105-0.0124 nM), respectively, comparable to the parent Dd2 EC_50_= 0.0128 nM (0.0124-0.0132 nM) (Fig S1) [21]. Altogether, our results indicate that the mechanism-of-action of MMV1580853 is distinct from that of chloroquine or artemisinin.

**Figure 4.**
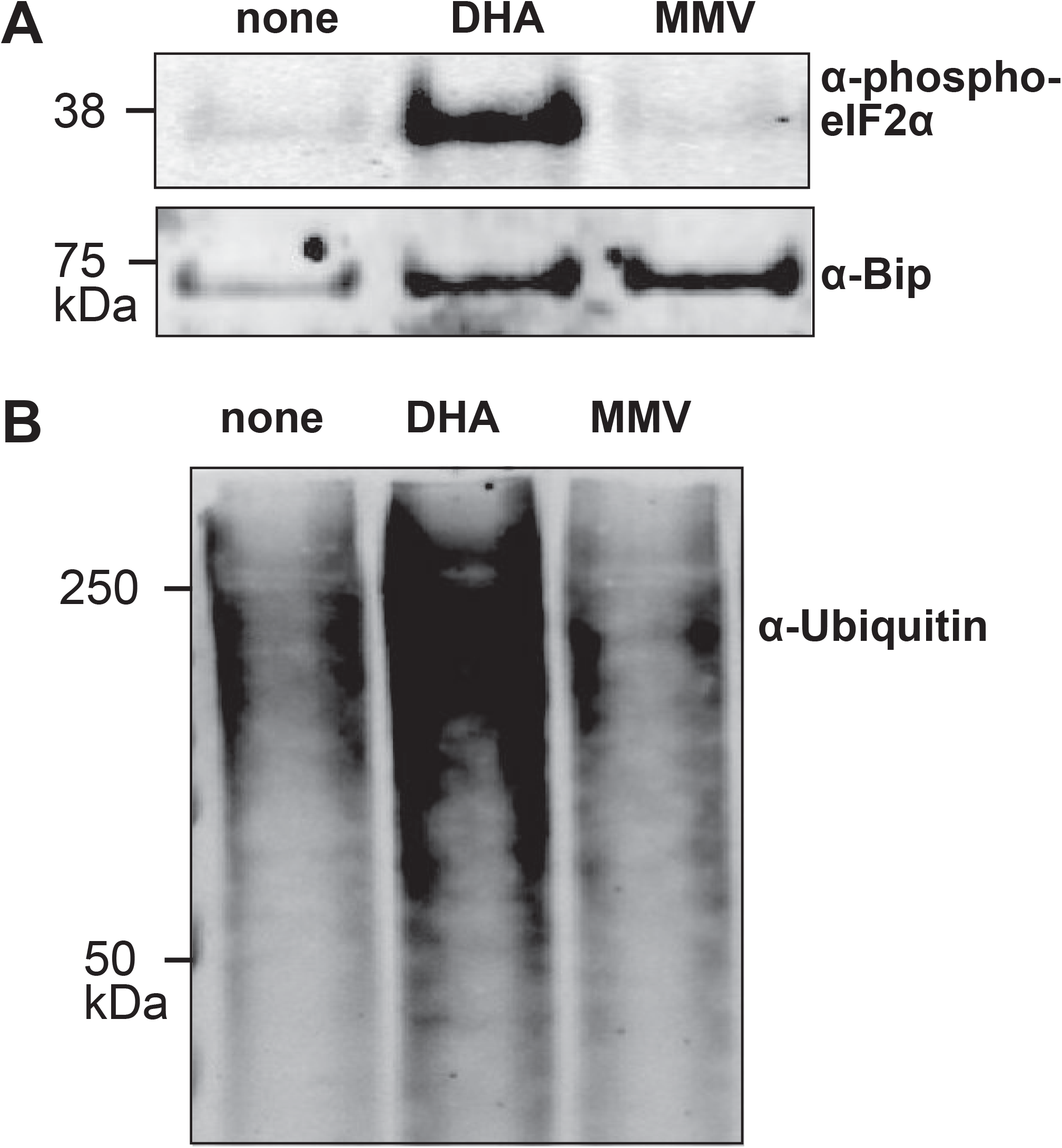
MMV1580853 treatment does not phenocopy artemisinin inhibition. A. Accumulation of polyubiquitinated proteins in inhibitor-treated *P. falciparum* parasites by Western blot. B. Accumulation of phosphorylated eIF2α in inhibitor-treated *P. falciparum* parasites by Western blot. Early trophozoite parasites (3D7) were grown for 2h in the presence or absence of indicated drugs at 3xEC_50_. Abbreviations as follows: none, no drug; DHA, dihydroartemisinin; MMV, MMV1580853. Bip serves as a loading control.

To identify the molecular target of MMV1580853, we attempted to select for resistant parasites *in vitro* in W2, Dd2, and 3D7 clonal parental strains. Parasites were treated with sublethal doses of ethyl methanesulfonate to increase the mutation rate and selected with either a lethal dose of MMV1580853 for up to 60 days or stepwise drug pressure starting at 1x EC_50_ and increasing over time [22]. Despite these different approaches to increase the likelihood of selecting resistant parasites, no MMV1580853-resistant parasites were recovered. *In vitro* resistance selection has been shown to be more successful on slow- and medium-acting compounds with fast-acting compounds often being “irresistible” [23,24].

Overall, MMV1580853 boasts a very fast killing rate comparable to dihydroartemisinin, indicating the potential to match the remarkable clinical efficacy of ARTs. Its mechanism-of-action is distinct from known fast-acting inhibitors, chloroquine and artemisinin, and it does not show cross resistance with artemisinin-resistant strains. Future directions include compound development and further investigation of its mechanism-of-action. To aid medicinal chemistry optimization, compounds in Pandemic Response Box library were tested in a number of ADME assays, in which MMV1580853 showed low mammalian cell cytotoxicity [25,26]. Moreover, MMV1580853 has been reported to have activity against transmission stages of *P. falciparum* and showed a selectivity index of >100 against HepG2 cells and >1000 against CHO cells [27]. *Pf*ATP4 inhibitors are another class of fast-acting antimalarials in pre-clinical development and clinical trials [28,29]. It will be valuable to test MMV1580853 for known *Pf*ATP4 inhibition phenotypes and against *Pf*ATP4 resistant mutants to determine whether their mechanisms-of-action are overlapping or distinct. MMV1580853 is a valuable addition to the portfolio of compounds under investigation to replace artemisinins.

## MATERIALS AND METHODS

### Parasite growth assays and transfections

*Plasmodium falciparum* parasites were grown in human erythrocytes obtained from the Stanford Blood Center at 2% hematocrit in RPMI 1640 medium (Gibco). Medium was supplemented with 0.25% Albumax II (Gibco), 2 g/liter sodium bicarbonate (Fisher), 0.1mM hypoxanthine (Sigma), 25mM HEPES pH 7.4 (Sigma), and 50 ug/liter gentamicin (Gold Biotechnology). Parasites were incubated at 37°C, 5% O_2_ and 5% CO_2_. *Plasmodium falciparum* W2 (MRA-157), Dd2 (MRA-150), 3D7 (MRA-102), were obtained from MRA (now BEI). NF54^Cas 9+T7 polymerase^ was kindly provided by Jacquin Niles [13].

To measure growth inhibition, *P. falciparum* cultures were grown in 96-well round-bottom plates containing serial dilutions of drugs. All assays were initiated with sorbitol synchronized ring-stage parasites at 1% parasitemia and 0.5% hematocrit in a final volume of 125 μL. For EC_50_ determination, parasites were treated continuously for 72 hr. For measurement of parasite viability (rate of kill), parasites were dosed at 10xEC_50_ drug for 1-72h, washed to remove drug, and cultured until the parasites reached the trophozoite stage of the following cycle (72 h from assay start). All assays were terminated by fixing with 1% paraformaldehyde and stained with 50nM YOYO-1. Parasitemia was determined by flow cytometry. Data was analyzed with BD C6 Accuri C-Sampler software and fitted to a sigmoidal dose-inhibition curve by GraphPad Prism.

For generation of knockdown strains, transfections were performed with NF54^Cas 9+T7 polymerase^ parasites. 100 μg of each plasmid was ethanol precipitated and resuspended with 30μL of sterile TE buffer, 170μL of cytomix, and 200μL of packed erythrocytes into a 0.2 cm electroporation cuvettes. Erythrocytes were electroporated at 310 V, 950μF, infinite resistance in a Gene Pulser Xcell electroporation system (Bio-Rad). Immediately after, schizont-stage parasites were added and allowed to invade in the presence of 500nM anhydrotetracycline (ATc) (Sigma). 2.5μg/ml Blasticidin S (Research Products International) was applied 3 days post-transfection. Cultures were maintained for up to 30 days to select for transfected parasites. To determine growth time course of TetR/DOZI strains, ring-stage parasites were washed 2x in growth media to remove ATc. Parasites were separated into cultures treated with 0-500nM ATc. Samples were collected at the schizont stage in each growth cycle for flow cytometry analysis. Parasites in each condition were diluted equally every growth cycle for up to five growth cycles. For parasitemia measurements, parasite-infected or uninfected RBCs were incubated with live-cell DNA stain dihydroethidium (ThermoFisher D23107) for 30 minutes at a dilution of 1:300 (5mM stock solution). Parasites were analyzed on a BD Accuri C6 flow cytometer and 100,000 events were recorded.

### Cloning and vector construction

Oligonucleotides and gBlocks were ordered from IDT. Molecular cloning was performed using Gibson Assembly (NEB) or In-Fusion Cloning (Clontech). For CRISPR-Cas9-based editing of endogenous PF3D7_1128400 (GGPPS), PF3D7_0826400 (CPT), and PF3D7_0202700 (OPP) single guide RNAs (sgRNAs) were designed using the CRISPR/Cas9 guide sequence program CRISPR. To generate a linear plasmid for CRISPR-Cas9-based editing, left and right homology regions, LHR and RHR respectively, were amplified for each gene. A gBlock containing the recoded C-terminal sequence after the CRISPR cut site and a triple hemagglutinin (HA) tag was synthesized with appropriate overhangs for Gibson Assembly. This fragment and the LHR were simultaneously cloned into the FseI/ApaI site of the linear plasmid pSN054-V5. Next, the appropriate RHR and a gBlock containing the sgRNA expression cassette were simultaneously cloned into the AscI/I-SceI sites of the resultant vectors to generate each TetR-DOZI plasmid. The sequences used for each construct were as follows:

#### PF3D7_1128400 (GGPPS)

*sgRNA*: ATATGTCTTGAAATACGTAA

*LHR*:TATATCATTGGACAATTTATTATACAAAAAGGTCGAAAATATAGCAATTCTTATGGGGGAATATTTTCAAGTTATTATTTAAAAAATATATATATATTTTTTTTTATTTTATTTATTTATTTATTTCATTTTTATATTATCTTTATGAATAAATATATACATATAATATAATTTTATTTTTATTAGGTCCATGATGATTATATAGATACCTTTGGAGATTCTAAAAAGACGGGAAAAGTTGGCTCAGATATTCAAAATAATAAATTAACGTGGCCCTTGATAAAAGTATCTTTACAAGAAATATTACATAGACATATATATATTTATATGTATTCATTTTATTTTATATTATATTATATTATATTTTATTATTATTATTATTTTTTTTTTTTTGTAGGCATTTGAACTATGTTCACAACCTGAAAAAGAGGACATAATAAGAAATTATGGGAAAGATAATGTAACATGTATTAAGTTTATTAATGATATATATGAACATTATAATATCAGGGATCATTATGTGGAATATGAAAAGAAGCAGAAGATGAAAATATTAGAGTAATAATATATAAGCATAAGTATACATGAATATATAAATATAAATATATATATATATATATATATATATATATATATATTTTTTTTTTTTTTTTTTTTTTTTTTTTTTGTAGAGCCATAAACCAATTGCAT CATGAAGGTAACATATAAATAATTTAATTTTTTTTTTTAATTATTTATTTATACATACATATATATATTATATATATATCAAAAAATTTTATGTATTAATATATTTATATCTTCGTTTTTTTTTTTCGTAGGTATAGA

*RHR*:GAGATAACAACCTTAACAACTTTTGTTTAAATATGATCAAGTTTAAAAAAAAAAAAAAATAATAATAATAAAACATTGAATGTAGATACATTATTAGTTTATATGTATATTTTTTTTTTTTTTTTTTTTTTTTTTTTTCTTATTCATTATTATATATATATATATATATGTATATATATTTGTGTACATCATTATATTCCTTTTTTTAAACCAGTGTTGTTATTTATCTTATTTTATTTCCTTATTTTTATTATTACTTTATTTATTTTTTTCTTTTTTTGTGGATCTCACATTTTACAATATAAAGATATTTTATTTTGAAAAATTTTTTACTACCATAGAAACATTTTAAAAATGATATTATGAATAATGAAATATTGTGTATATACTAGTTCTATATATATATATATATATATATATATATATTTATATATTTTATATGTACCTTTTAAAACGTCCAATATCCATATAATATAAATTTTTTCCCTCATAAAATATATATATATATATATGTATAGTTTTATTTATTTTATCCCTTCAATATCTTTAAATACTTTTTTTTTTTTTACAACATATCGTATTAATATCTAATCAAATATATTTATATGTATATATATATATATATATATTTCATTTCATTTCATTTATTCTTATTAAAATATGGATATCGAGAATTCAAAAAAACACCTAAGCGAATTACAAGAAGAAAGAAGAAGACACAAAATTACACTTCCCGATATGTTTAAAAGGAAAGTTCTTATAAAGGAAGAAAATAATGATACTTATGAAAAGAATTTAGAGACAAAAAATGATATGTGTTATTCATTGAATAGATCAGATGATGAGAAGG

*C-terminal gBlock*: GTACGTCCTGAAATACGTAATGGACATCCTGTTCACAGGCGCT

#### PF3D7_0826400 (CPT)

*sgRNA*: CGAGTATGAAAATATCCACT

*LHR*:CCCACACAAGAAGAAGAACTAAAATATCCTGATCATGAAAATGAAAATGAATTAAAATTTGATGGAAAATGTCTTTGCAGGGAAAAAGTAAAATACAATGAAGAGCAATTAGAAATGTAAGTATATATATATGTATATATTTGGAGAATCATTTATTATTTAATATTTAATTTTTTTTTTTTTAATTTATTTTTTTTTAATTTCTATTTATTCTTTTACATCTTTTATTTCATTTCATTTTTCTTTTTTTGCAGCGTTAATTATCATAATAAATTGCTTACAAGTGATTTGCCTCCCCCTAATATTTTAATAAGAACATCAGGTGAACAAAGACTATCCGATTTTATGTTATATCAGGTAAATATATATATATAAATATATATATGCCGTTATGATATATAAAAATATATATTTCAAACAAATAATATATCTATATAAACGTTTATAAATTTTTATTCATATATATATATATATATATATATATACATATATTCCCCTTTTCAGATATCAGAGTTTACCGAAATTTATTTTATTAATGAATATTGGCCAGTATTTAATTTTTTACAATTTATTTATATAATATTACATTATACCCTTTTTCAAACAA

*RHR*:AATTATATATATAAATAAATATAAAGACATATATATATATATATATATATATATATATATATATTTATATATATTTATTTATTTGTTTATATTTTTTTTTTTTTTTTTTTTTTTTTTTTGTGTTATTGTTTTATTTGTTTTTTTTTTGTGTATTTGTAAATTAATTTGTTATTTTTATTCTCTTATTTCTTTAAATGAAAATATAAAAACGTGTAAATATTTTGAAATCATCAGTTTTGATATTTCTAAATGAAAATCATGCATTTTTATGTTAAAATAAAAGTTCACCTGTACACACATATATATATGTACATATTTTGAAAGATACAGAAACGTAAAAATAACAAAAAATGAAAAATAAAAAAAAAATAAAAAAAATGTATATAATGTAATGAAATGAGATAAATGATTTATAATTATTAATTATTTAGTAATCCAAAAGTATCTCATATAAACAAATATATATATATATATATATATACATATATATGTTTATATTTTCTTTTTATAATACGGTGGGAGGCTCCATATATACACATGGGACATAGTCCTGCAAATGATTATATGTAAAATTAGGGCCCAAGTTAATTTTAGCTGATGCTCCC

*C-terminal gBlock*: CGAAGTGGATCTTCTCCTACTCGAATCCTTGTCCACAC

#### PF3D7_0202700 (OPP)

*sgRNA*: TGATATAAAACAAAGTAGCG

*LHR*:CTCTTAGCACGTGCTAGTTCTATTTTTGCTGGTACGGGTTCACCAAAAATTTGTAGAAGTTTCTCTTATGTTGTCGAAAGTTTAATAAAAGGAGAATTCTTACAAAGAAATTTAAAATTTAATAATGTTGAAGAAGCACTCAAAATGTATTTAATTAAATCATATCATAAAACAGCTTCTCTTTTTTCTCATTTATTTGCATGTATAGCCATTCTATCATTTAAAAATGATACTATTATACAATTATGTTTTAATTTAGGATTACACATAGGTATGGCTTTTCAATTATATGATGATTATCTAGATTATAAAATCGATGACAATACAAACAAACCTATATTAAATGATTTAAAAAATAATATTAAAACAGCTCCCTTATTATTTTCCTATAATTATAACCCTCAAGTTATCTTACAATTAATTAATAAAAATTCATATACAAATAATGATATTGAAAATATTTTATATTATATCCAACATTCTAATAGTATGAAAAAAAATGAATTGTGTTCATTATTACATATAAAAAAGGCATCTGATATTCTATACTCCTTAATATCTCATTGTAATAAACCTAGTACAAATAAAAACAATACCAAACATGA

*RHR*:TAAAAGAATAAATAAAACACATATATATATATATATATATATATATATATATTAATAATTAATATAAAAGATTATTTTAATTTTTATTCAATAACATATACAATATCAAACATATATATTATAATATTATTAAACATCTTCAATATTGTATTATTTAAAAAACTTAATAATATATACATATATTTTATATATATATATATTTTTGTTAATTAGTTTGAAAAAAATAAAAAACAATTTATAATATTTAATCCATTTATTTATATGTTTCATTAATTTATTGTTACTACATATTTGTTATATATATTATTTTTTTGTCTTTTTAACACTTTTCATATTATTTGTATAAAAAAAATTTCCATTATTTCATATACACCCCACATTCCTTTTATCTTTTTTAAATTTTATTTATCTCATGTATTATTTGTGCCCCTTACAACTATAATAAAATAAATATAAACAAATAAATAAATACTAAAGAAAAAAAAAAAAAAAAAAAAAATTCTTTTTTTTTTAAATAAAAATATATTTTTTTTTTTTTTTTTACCAAATATTAAATTTAGTACAATTCCTTTTACTCTCTTTCCAAAATACATTTTCCTCCTC

*C-terminal gBlock*:TGATATCAAGCAGAGCAGTGAAGCATTAATTAATTTAATCTTAAACGTGTTATCAAGAAACGTCAAA

### Parasite Imaging

Synchronized early ring-stage 3D7 parasites were treated with either no drug, with 100nM of MMV1580853, Pyrimethamine, or Chloroquine. Parasite morphology was monitored every 6 hours for 48 hours. Cultured erythrocytes were spread thinly onto a glass slide and fixed with methanol. The slides were incubated with giemsa solution for 10 minutes. Glass slides were washed with DI water and air dried. Cell morphologies were observed under a light microscope using a 100x oil immersion objective lens. Images were taken using a camera mounted to the microscope.

### Western blots

3D7 parasites were treated as mid-trophozoites (30 hr post invasion) for either 10 hr (hemoglobin), or 2 hr (ubiquitin and phospho-eIF2α) with either 100nM for MMV1580853, DHA, chloroquine, or no drug. Parasites were extracted from infected erythrocytes by lysing with 0.1% saponin for 5min on ice. Pellets were washed three times with cold PBS. Samples for ubiquitin analysis contained EDTA-free protease inhibitor cocktail (Roche) and 20mM N-ethylmaleimide (Sigma) in both the lysis and wash buffers. Ubiquitin and phospho-eIF2α samples were resuspended in 1X NuPAGE LDS sample buffer, then boiled for 15 min before running on a 4-12% Bis-Tris NuPAGE gel with MOPS running buffer. Hemoglobin samples were resuspended in 1X Novex Tris-Glycine SDS sample buffer then boiled for 15 min before running on a 16% Tris-Glycine Novex gel with Novex Tris-Glycine SDS running buffer. Gels were transferred onto nitrocellulose membranes using the Trans-Blot Turbo system (Bio-Rad). Membranes were blocked overnight at 4C with 5% nonfat dry milk and 3% BSA in TBST (100mM Tris-Cl, 0.5M NaCl, 0.05% Tween-20, pH 7,5). All antibodies were dilute in 1:1 mixture of blocking buffer and TBST. Blots were incubated with primary antibodies either for 1hr RT or overnight at 4C with 1:100 ubiquitin (cell signaling technology #58395, 1:1000 Phospho-eIF2α (cell signaling technology #9721), 1:10000 hemoglobin *β* (cell signaling technology #84934), *Pf*BIP (BEI resources #MRA-1246.Blots were subsequently washed 3x with TBST and incubated for 1hr at RT with 1:10,000 dilution of LI-COR IRDye secondary antibodies (800CW or 680RD). Blots were then washed 3x TBST and 1x PBS prior to being imaged on a Bio-Rad ChemiDoc gel imager. Protein levels for the hemoglobin blot were quantified using densitometric analysis in Fiji (ImageJ). Undigested hemoglobin band intensities were normalized relative to untreated parasite controls.

### Mutagenesis and resistance selection

Parental W2, Dd2, and 3D7 strains were sorted by RBC size on a Sony SH800S Cell Sorter, each well was given one RBC, only wells in which the RBC was infected grew up. Late-stage parental parasites were incubated with 0-1.5 mM ethyl methanesulfonate (EMS, 8 concentrations total) for 2 hrs and then washed 3x with growth media. Parasites were transferred to flasks containing 10mL of media, approximately 10% parasitemia, 2% hematocrit. Parasites were then exposed to a constant lethal dose of MMV1580853 (100nM) for either 15 days and allowed to recover for another 45 days or exposed for a full 60 days. Additionally, parasites were treated with sublethal drug dose at 1 x EC_50_ (15nM) and then drug concentration was slowly increased over time. Cultures were fed daily for the first week and then every 3 days thereafter with the inhibitor being replenished at every feeding. Once a week, cultures were split in half and fresh RBCs added.

## ACKNOWLEDGEMENTS

We thank Medicines for Malaria Venture (MMV) for providing the Pandemic Response compound library. This work was supported by the following funding: NIH 1R01AI141366 (EY) and NIH 5T32HG000044 (SCK). EY is also a Chan Zuckerberg Biohub – San Francisco Investigator and supported by the Burroughs-Wellcome Fund.

## Supporting Information

**Figure S1.**
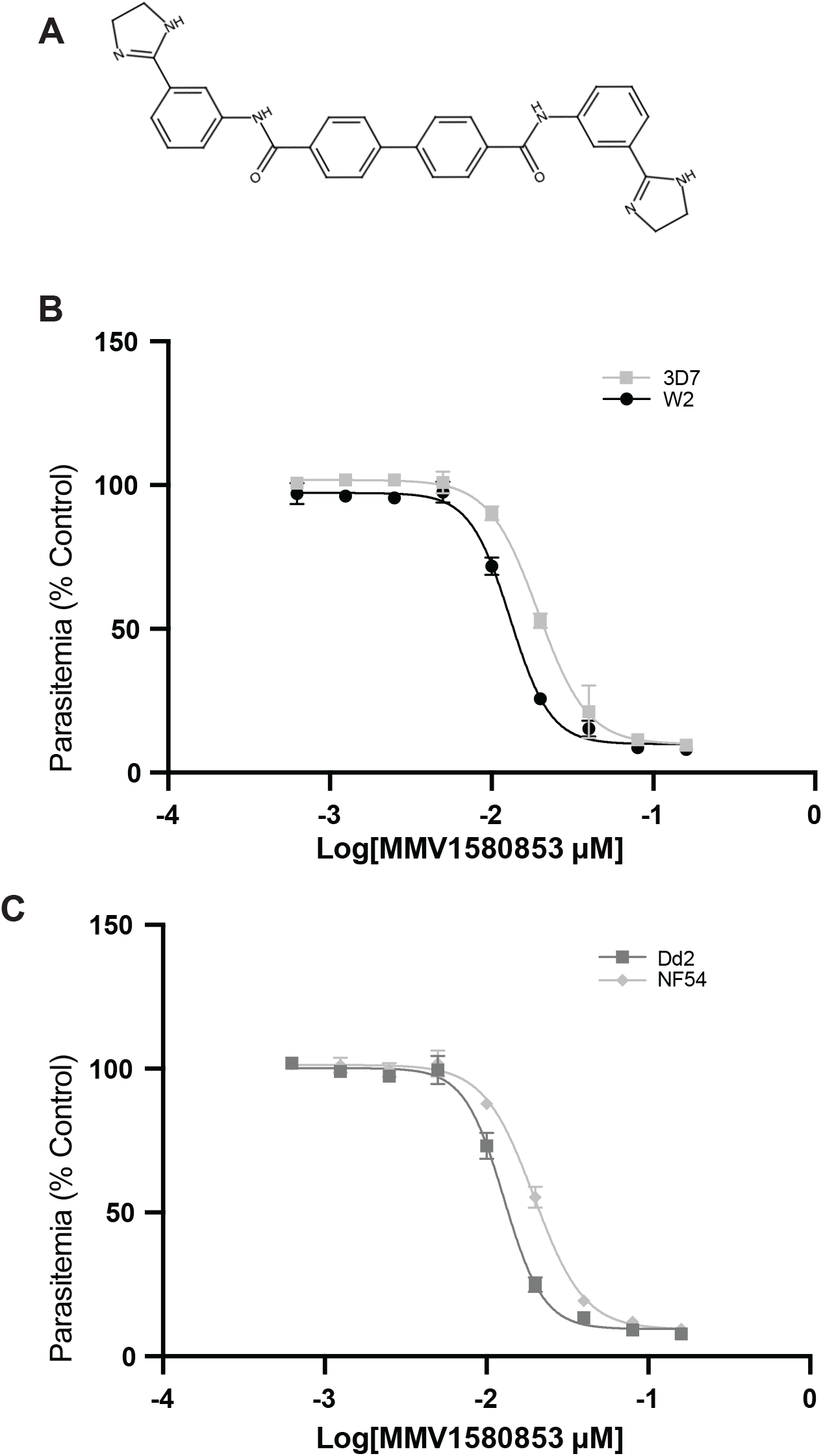
MMV1580853 inhibition of parasite growth. A. Structure of MMV1580853 B-C. 72h dose-dependent growth inhibition of blood-stage *P*.*falciparum* strains: (B) chloroquine-resistant W2 and chloroquine-susceptible 3D7; (C) parent Dd2 (background strain for Kelch13 mutants) and parent NF54^pCRISPR^ (background strain for TetDOZI KD strains). Results are the mean and 95% CI of three independent experiments after subtraction of background fluorescence of uninfected RBCs. Growth is normalized relative to untreated controls.

## Notes

### Competing Interest Statement

The authors have declared no competing interest.

